# Expanding the Distribution and Phylogenetic Insights of *Chrysobrycon mojicai* in the Peruvian Amazon: Morphological and Molecular Analyses with Taxonomic Corrections

**DOI:** 10.1101/2024.08.02.606444

**Authors:** Junior Chuctaya, Morgan Ruiz-Tafur, Dario Faustino-Fuster, Vanessa Meza-Vargas, Carmen Garcia-Davila, Diana Castro-Ruiz, Carlos Angulo, James Anyelo Vanegas-Ríos

## Abstract

This study focuses on the genus *Chrysobrycon*, particularly *Chrysobrycon mojicai*, which was initially described in the Amacayacu National Natural Park in Colombia. Here, we document a new geographical record of *C. mojicai* in various locations of the Peruvian Amazon, including the Nanay, Putumayo, Tapiche, and Tigre Rivers basins. Based on morphological, morphometric, and molecular analyses, we confirm the presence of *C. mojicai* in these new locations, expanding its known distribution. Morphological features such as the distinct shape of the hypertrophied scales and the specific arrangement of teeth were used to confirm its identity. Molecular data, obtained through cytochrome oxidase I (COI) gene sequencing, provide additional validation and contribute to understanding its phylogenetic relationships within the Stevardiini tribe. Our phylogenetic analysis reveals unresolved relationships within the tribe, particularly in the genus *Gephyrocharax*, and highlights discrepancies in the current taxonomic framework, with *C. mojicai* showing close genetic affinity to *C. myersi* from the Pachitea River basin. The study also presents morphometric information of the holotype of C. mojicai, specifically the percentages of measurements relative to the head, which were not included in the original description. It also includes ecological observations of the habitats where *C. mojicai* was collected, noting its presence in blackwater and mixed water streams characterized by fluctuating water levels and specific physical and chemical parameters. Additionally, the study restricts the distribution of *C. guahibo* for Colombia and invalidates the COI sequence of *Hysteronotus megalostomus* available in molecular databases. This research not only expands the known distribution of *C. mojicai* but also underscores the need for further taxonomic and ecological studies to resolve existing ambiguities within the Stevardiini subfamily.

## Introduction

The family Characidae (Characiformes) is one of the most diverse groups in the Neotropics, currently encompassing 1,259 valid species, which 207 species (17 %) having been described in the last decade [1]. Historically, Characidae has been diagnosed by the absence of the supraorbital bone [2–5]. This family is widely distributed, ranging from southwest Texas and Mexico through Central America to Patagonia, Argentina in South America. Most species are found in the Amazon and Orinoco River basins, as well as in the Guianas rivers, with over 594 species identified in these regions, distributed across 98 genera [6]. Despite extensive research, systematic aspects of Characidae remain incomplete, including its species richness, morphological diversity, and the limits of intra- and intergeneric relationships, with many genera still considered paraphyletic and polyphyletic, as well as unresolved internal phylogenetic relationships.

The family Characidae is represented by eight subfamilies and several *incertae sedis* taxa [7]. Among these is the subfamily Stevardiinae, which includes 375 valid species distributed across 45 genera, accounting for 30 % of the family’s species [1]. Within Stevardiinae seven monophyletic tribes have been proposed, along with some unresolved clades [8]. One such tribes, Stevardiini, comprises the genera *Chrysobrycon* Weitzman & Menezes 1998, *Corynopoma* Gill 1858, *Gephyrocharax* Eigenmann 1912, *Pseudocorynopoma* Perugia 1891, *Pterobrycon* Eigenmann 1913, and the recently added *Hysteronotus* Eigenmann 1911 [9].

The Stevardiini form a well-supported monophyletic tribe consisting of two sister clades. The first clade includes *Chrysobrycon*, *Hysteronotus*, and *Pseudocorynopoma*, while the second clade is composed of *Corynopoma*, *Gephyrocharax*, and *Pterobrycon*. This tribe is supported by 30 DNA synapomorphies and three morphological traits: (1) a well-developed groove above the eyes, (2) an inner premaxillary tooth row with a maximum of five cusps, and (3) the dorsal fin origin posterior to the vertical line passing through the origin of the anal fin [9]. The monophyly of Stevardiini has been corroborated by multiple studies [7, 9, 10].

The genus *Chrysobrycon*, currently includes the following species: *C. hesperus* (Böhlke 1958), described from a tributary of the Napo River basin; *C. myersi* (Weitzman & Thomerson 1970), described from a tributary of the Pachitea River; *C. eliasi* Vanegas-Ríos, Azpelicueta & Ortega 2011, described from the Madre de Dios River basin; *C. yoliae* Vanegas-Ríos, Azpelicueta & Ortega 2014, described from the Abujau River, a tributary of the Ucayali River; *C. guahibo* Vanegas-Ríos, Urbano-Bonilla & Azpelicueta 2015, described from the Guaviare River in the Orinoco basin; *C. mojicai* Vanegas-Ríos & Urbano-Bonilla 2017, described from the Matamatá River, a tributary of the Amazon River in Colombia; and *C. calamar* Vanegas-Ríos, Urbano- Bonilla & Sánchez-Garcés 2024, described from the upper portion of the Vaupés basin in Colombia. These species illustrate the diversity within *Chrysobrycon*, each being described from distinct geographical locations. This distribution highlights the varied ecosystems they inhabit and underscores the genus is ecological diversity.

*Chrysobrycon* is characterized by several distinctive features, including hypertrophied scales on the lower of the caudal-fin lobe that forms a laterally open pocket in adult males [11–14]. This pocket is primarily composed of a specialized pocket scale, which is relatively small compared to those in other related stevardiins. This scale is somewhat elongated, curved, confined to the dorsal region of the pocket opening, and horizontally folded, giving its lateral face a concave shape [11]. Another key characteristic of *Chrysobrycon* is the extensive contact of the frontals along the midline [15]. Additional distinctive characteristics of the genus include a superior mouth, a dorsal fin located posterior to the origin of the anal fin, insemination, and the presence of numerous bony hooks on the pelvic, anal, and caudal fins in adult males [11, 16, 17].

In Peru, five species of *Chrysobrycon* have been documented; however, only four have been recorded within the Loreto region [18]. These species are *C. eliasi*, *C. hesperus*, *C. myersi*, and *C. guahibo,* the latter of which was originally described from the Orinoco River basin. The occurrence of *C. guahibo* in the Amazon basin in Loreto region necessitates further investigation to better understand its distribution. This finding adds a new dimension to our understanding of the geographical range of *C. guahibo*.

*Chrysobrycon mojicai* was initially described by Vanegas-Ríos & Urbano- Bonilla [15] from a tributary of the Matamatá River in the Amacayacu National Natural Park in Colombia. To date, no additional geographic records of this species have been reported from other aquatic environments. Consequently, our objective is to corroborate the presence of *C. mojicai* in the Peruvian Amazon basin through a morphological comparison. Furthermore, we aim to assess the potential placement of *C. mojicai* using cytochrome oxidase subunit I (*coI*) data, which are for first time presented for the species, as part of a phylogenetic comparison including other members of Stevardiini.

## Material and Methods

### Ethical Statement

Fish sampling was conducted in the Loreto region of Peru, under collection permits DIRECTORAL RESOLUTION N° 00352-2024-PRODUCE/DGPCHDI, issued by the Ministry of Production of Peru (PRODUCE). This work was part of the project titled "Assessment of fish diversity in streams along the Iquitos-Nauta Road axis, Loreto, Peru. Additionally, part of the samples was collected under the collection permit DIRECTORAL RESOLUTION No. 00495-2024-PRODUCE/DGPCHDI, within the project “Study of type species described in Peru focusing on an integrative analysis for the strengthening and validation of the fish DNA barcoding database for use in environmental DNA (eDNA)”, funded by CONCYTEC-PROCIENCIA. No surgical or experimental procedures were performed on live fish.

### Specimen Collection

Fish sampling was carried out using shore seine nets (8 meters long and 2 meters high) with a mesh size of 3.0 mm. A portable multiparameter meter, HANNA model HI98194, was employed to record environmental parameters. Collected specimens were anesthetized with eugenol diluted in 96% alcohol [19]. From molecular analysis, 1 cm^3^ of muscle tissue was extracted from a selected specimen and preserved in 96% alcohol. Additional photographs of live specimens were taken using a Nikon D3100 camera.

Following photography, specimens were fixed in 10% formalin, rinsed in water, and subsequently preserved in 70% ethyl alcohol. The specimens studied are deposited in the Ichthyological Collection of the Research Institute of the PeruvianAmazon (CIIAP) and Natural History Museum of the San Marcos University (MUSM).

### Morphological Analyses

Morphometric and meristic data were collected following the methodology described by Vanegas-Ríos & Urbano-Bonilla [15]. Measurements were taken using an IGAGING digital caliper with a precision of 0.01 mm and are presented as percentages of the standard length (LS) and head length (HL). The identification of the specimens was based on the description of *C. mojicai* [15] and additional comparative specimens. These comparative specimens are deposited in institutions, which acronyms follow Sabaj [20].

To analyze shape variability of the studied specimens in comparison with the type series of *C. mojicai,* we performed a principal component analysis based on the covariance matrix obtained from the size-corrected measurements using the Burnaby’s procedure [21]. Missing values in the morphometric data were imputed using the EM algorithm (100 iterations) [22, 23]. These procedures were conducted using PAST 4.12 [24], IBM SPSS Statistics 26.0 [25], and GraphPad Prism 9.4.1(GraphPad Software, San Diego, CA, USA). The geographic distribution map was prepared using Quantum GIS 2.18.10 software [28].

### DNA extraction, mitochondrial COI amplification and sequencing

DNA extraction was performed a standard salt extraction protocol [26], starting with 50 mg of muscle tissue. DNA amplification was conducted with primers L5698-Asn (5’ AGG CCT CGA TCC TAC AAA GKT TTA GTT AAC 3’) and FishR1 (5’ TAG ACT TCT GGG TGG CCA AAG AAT CA 3’), as designed by Hubert et al. [27]. The amplification reaction was carried out in a total volume of 10 μl, containing 0.7 μl of Taq polymerase (1 U/μl), 1.0 μl of template DNA (100 ng/μl), 1.0 μl of Buffer 10X, 1.7 μl of dNTPs (2 mM), 1.0 μl of MgSO4 (25 mM), 0.5 μl of each primer (10 μM), and 3.6 μl of ultrapure water. The thermal cycling conditions were as follows: initial denaturation at 94 °C for 2 min, followed by 35 cycles of denaturation at 94 °C for 30 s, annealing at 50 °C for 40 s, and extension at 72 °C for 1 min, followed by a final extension at 72 °C for 10 min. The amplified products were separated by electrophoresis on 2 % agarose gels. Sequencing was performed on a 3500XL Genetic Analyzer (Applied Biosystems) using the BigDyeTM Tr v3.1 Cycle Sequencing kit following the manufacturer’s instructions.

### Molecular analysis

DNA chromatograms were edited using BioEdit [29]. The obtained *coI* sequences were verified using the BLAST search engine provided by the National Center for Biotechnology Information (NCBI), a primary public repository for DNA barcode sequences. Sequences were aligned using Muscle with default setting as implemented in MEGA 11[30]. The Kimura 2-parameter model (K2P) [31] was employed to calculate pairwise distances between specimens collected in this study and several other specimens of the same genus retrieved from NCBI (S1 Table 1). A total of 72 *coI* sequences were analyzed, of which 67 were retrieved from GenBank, including the two sequences of *Chrysobrycon mojicai* here obtained, which were deposited in the GenBank database with the codes PQ118467 - PQ118468. Of these sequences, 39 belonged to members of the Stevardinii tribe (genera *Chrysobrycon*, *Corynopoma*, *Gephyrocharax*, *Hysteronotus*, *Pseudocorynopoma*, and *Pterobrycon*). The remaining sequences, selected from related characids *sensu* Thomaz et al. [8], were treated as outgroups (*Glandulocauda melanogenys*, *G. caerulea* Menezes & Weitzman 2009, *G. melanopleura* (Ellis, 1911), *Eretmobrycon peruanus* (Müller & Troschel 1845), *Roeboides descalvadensis* Fowler 1932, *Tyttocharax tambopatensis* Weitzman & Ortega 1995, [32–37] and Cheirodontinae sequencies of Chuctaya et al. [38] and Javonillo et al. [39]. A phylogenetic analysis using maximum likelihood (ML) was run in RAxML-HPC2 on XSEDE 8.2.9 [40, 41] via CIPRES portal v3.3 [42] using the *coI* dataset. RAxML searches were performed with 10 parallel runs, starting with a randomly generated tree. Branch support was assessed using the rapid bootstrap algorithm with 1000 replicates. The optimal tree obtained was visualized using FigTree v1.4.3 [43].

**Table 1.** Meristic data of studied specimens of *Chrysobrycon mojicai* from the Nanay and Putumayo River basins (CIIAP-2315, 2324, 2613, and 2626).

|  | Holotype | Nanay River Basin |  |  | Putumayo River Basin |  |  |
| --- | --- | --- | --- | --- | --- | --- | --- |
|  | IAvH-P |  |  |  |  |  |  |
| Counts | 13932 | n | Min | Max | n | Min | Max |
| Unbranched rays anal fin | 4 | 4 | 5 | 6 | 5 | 4 | 5 |
| Branched rays of the anal fin | 30 | 4 | 24 | 27 | 5 | 25 | 27 |
| Number of dorsal-fin rays | 10 | 4 | 10 | 11 | 5 | 10 | 11 |
| Number of pelvic-fin rays | 8 | 4 | 8 | 8 | 5 | 8 | 8 |
| Number of pectoral-fin rays | 11 | 4 | 10 | 11 | 5 | 10 | 11 |
| Scales of lateral line | 45 | 4 | 42 | 45 | 5 | 44 | 47 |
| Scales of lateral line-dorsal fin origin | 7 | 4 | 6 | 7 | 5 | 6 | 6 |
| Scales of lateral line-pelvic fin origin | 5 | 4 | 4 | 5 | 5 | 4 | 5 |
| Predorsal scales | 21 | 4 | 19 | 27 | 5 | 21 | 27 |
| Scale rows around caudal peduncle | 15 | 4 | 15 | 16 | 5 | 15 | 16 |
| Premaxilla, outer row | 5 | 4 | 5 | 5 | 5 | 5 | 5 |
| Premaxilla, inner row | 5 | 4 | 4 | 4 | 5 | 4 | 4 |
| Premaxilla, inner row cusps | 5 | 4 | 5 | 5 | 5 | 5 | 5 |
| Maxillary teeth | 14 | 4 | 12 | 15 | 5 | 12 | 18 |
| Dentary teeth | 24 | 4 | 15 | 18 | 5 | 16 | 20 |

## Results

### *Chrysobrycon* Weitzman & Menezes 1998

*Chrysobrycon mojicai* Vanegas-Ríos & Urbano-Bonilla 2017 New record. **All from Peru: Loreto Department: Nanay River:** CIIAP-2613, 1, 39.79 mm SL, San Juan Baustista district, Tanshi creek, 3°54’33.85"S 73°24’37.65"W, 21 Nov. 2023, Ruiz M., Agurto E., Chuctaya J. CIIAP-2626, 4, 28.8–32.78 mm SL, San Juan Baustista district, Paujil creek, 3°57’53.56"S 73°25’2.26"W, 22 Nov. 2023, Ruiz M., Agurto E., Chuctaya J. MUSM 64997, 13, 30.76–38.58 mm SL, Alto Nanay district, 2°48’17.57"S 74°49’27.50"W, Aug. 2006, Hidalgo M. & Willink P. MUSM 65021, 1, 60.28 mm SL, Alto Nanay district, 2°47’29.00"S 74°49’36.84"W, 22 Aug. 2006, Hidalgo M. & Willink P. MUSM 65069, 3, 29.52–37.17 mm SL, Alto Nanay district, 2°48’39.67"S 74°45’17.57"W, 24 Aug. 2006, Hidalgo M. & Willink P. **Putumayo River:** CIIAP-2315, 4, 44.31–47.67 mm SL, Rosa Panduro district, Santa Elena creek, 01°50′00.7"S 73°18′32.1"W, 16 Apr. 2023, Ruiz, M., Sindy, J. CIIAP-2324, 1, 43.49 mm SL, Rosa Panduro district, Santa Elena creek, 01°49′52.4"S 073°18′19.8"W, 16 Apr. 2023, Ruiz, M., Sindy, J. MUSM 65549, 1, 43.77 mm SL, Rosa Panduro district, Ere River, 1°40’21.90"S 73°42’45.79"W, 17 Oct. 2012, Quispe R., Maldonado J., Hidalgo M. MUSM 64720, 2, 36.78 40.15 mm SL, unnamed creek tributary to Campuya River, 1°32’28.00"S 73°49’18.98"W, 28 Oct. 2012, Quispe R., Maldonado J. **Tapiche River:** MUSM 65367, 1, 41.68 mm SL, Tapiche district, Yanayacu River, 6°16’7.00"S 73°54’17.00"W, 18 Oct. 2014, Hidalgo M., Aldea M., Corahua I. MUSM 64353, 1, 43.11 mm SL, unnamed creek tributary to Tapiche River, Requena district, 5°10’57.55"S 73°52’1.90"W, 19 Aug. 2005, Hidalgo M. & Pezzi J. MUSM 65197, 1, 38.13 mm SL, **Tigre River:** MUSM 22920, 1, 41.18 mm SL, Urarinas district, Anguilayacu creek tributary to Corrientes River, 3°55’5.47"S 74°58’57.48"W, 19 Apr. 2004, Rengifo B. MUSM 23170, 6, 33.85–48.22 mm SL, Trompeteros district, Platanoyacu River, tributary to Corrientes River, 3°7’16.83"S, 76°12’39.10"W, 07 Oct. 2004, Martinez S. MUSM 26711, 2, 38.03–44.61 mm SL, Tigre district, Pucacuro creek tributary to Corrientes River, 3°26’2.69"S 75°25’21.51"W, 21 Oct. 2005, Gomez M. MUSM 26740, 1, 40.49 mm SL, Urarinas district, Brujulayacu creek tributary to Corrientes River, 3°55’14.98"S 74°58’43.96"W, 25 Oct. 2005, Gomez M. MUSM 69206, 13, 27.62–41.36 mm SL, 2°16’44.60"S 75°53’23.13"W, 11 Jul. 2019, Faustino N.

### Diagnosis and morphological description Species identity

The morphological, morphometric, and molecular analysis were conducted on individuals collected from Nanay and Putumayo basin. To determine the distribution area of *Chrysobrycon mojicai* in the Peruvian Amazon, individuals collected from the Tapiche and Tigre basins were included. The size of the studied specimens from the Nanay River basin was 28.8–60.3 mm SL, from the Putumayo River basin ranged from 43.5–47.7 mm SL, from the Tapiche River ranged from 41.7–43.1 mm SL and from the Tigre River ranged from 27.6– 48.2 mm SL. Only female specimens were observed in the Nanay river basin.

These specimens collected from Nanay and Putumayo basin (Fig 1) are characterized by the following characters: Scales cycloid. Lateral line completely pored: 42 (1), 44 (2), 45* (3), 46 (1), or 47 (2) scales. Anal-fin rays iv* (3), v (5), vi (1); 24 (1), 25 (3), 26 (2), or 27 (2). Pectoral-fin rays i (9), nine (7) or 10* (2). Pelvic-fin rays i(9), seven* (9) . Premaxillary teeth arranged in two rows, outer row with five (9) teeth, and inner row with four (9) teeth (inner row with five teeth in the holotype) (Table 1). The morphometric measurements fell within the ranges of the type specimens of *C. mojicai* (Table 2, see also [15]: Table 1), with a slight variation observed in upper jaw length (36.7–45.0 % HL *vs*. 43.1–49.8 % HL), snout to dorsal-fin origin (59.7–65.9 % SL *vs.* 62.9–70.5 % SL), and depth at dorsal-fin origin (24.1–29.1 % SL *vs*. 25.9–36.7).

**Fig 1.**
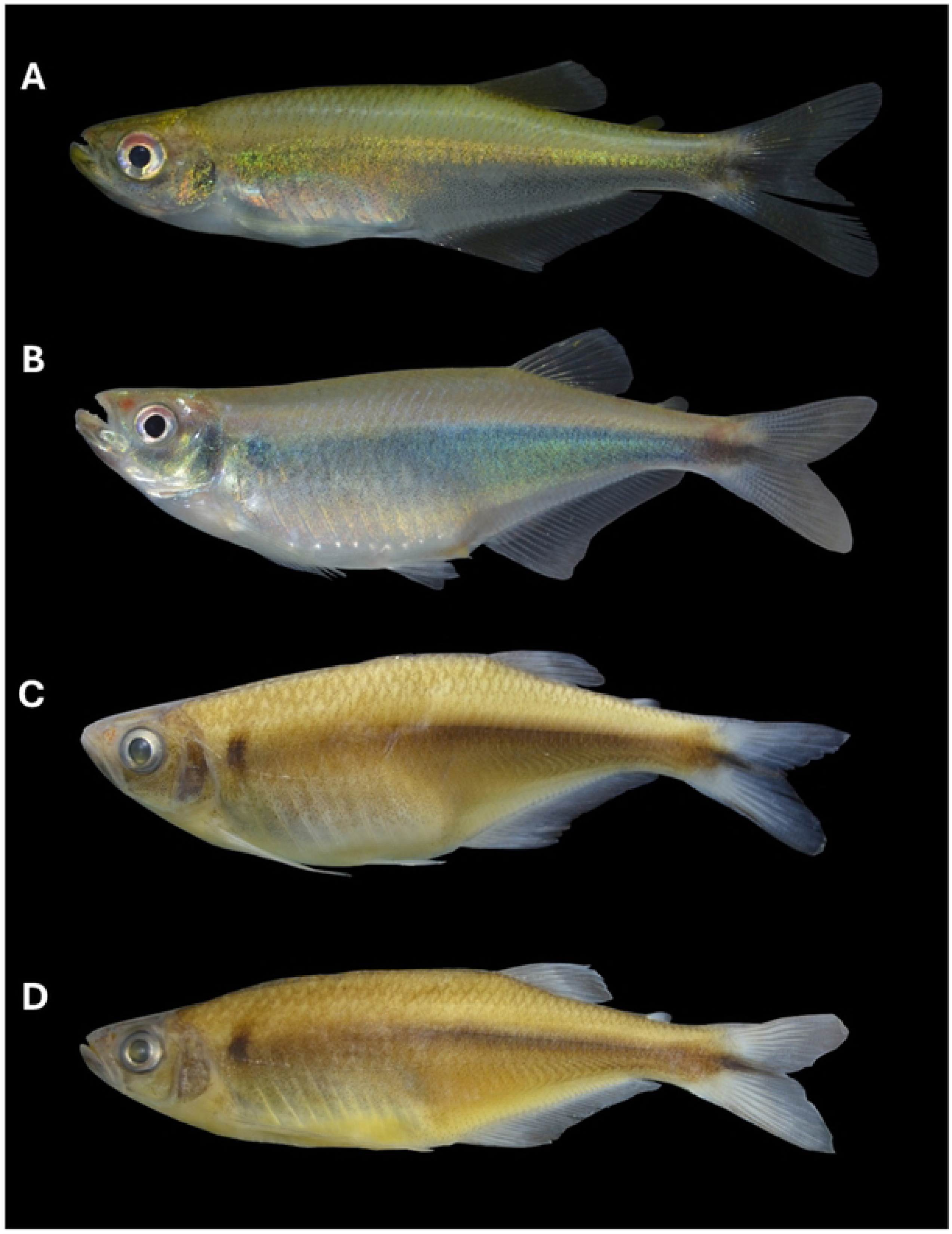
*Chrysobrycon mojicai* collected in the Peruvian Amazonian basin. (A) Photograph of live specimen from the Nanay River, female, CIIAP 2626, 32.78 mm SL. (B) Photograph of live specimen from the Putumayo River, female, CIIAP 2315, 45.89 mm SL. (C) Preserved specimen from Putumayo River, male, CIIAP 2315, 47.67 mm SL (D) Preserved specimen from Putumayo River, female, CIIAP 2315, 45.89 mm SL.

**Table 2.**
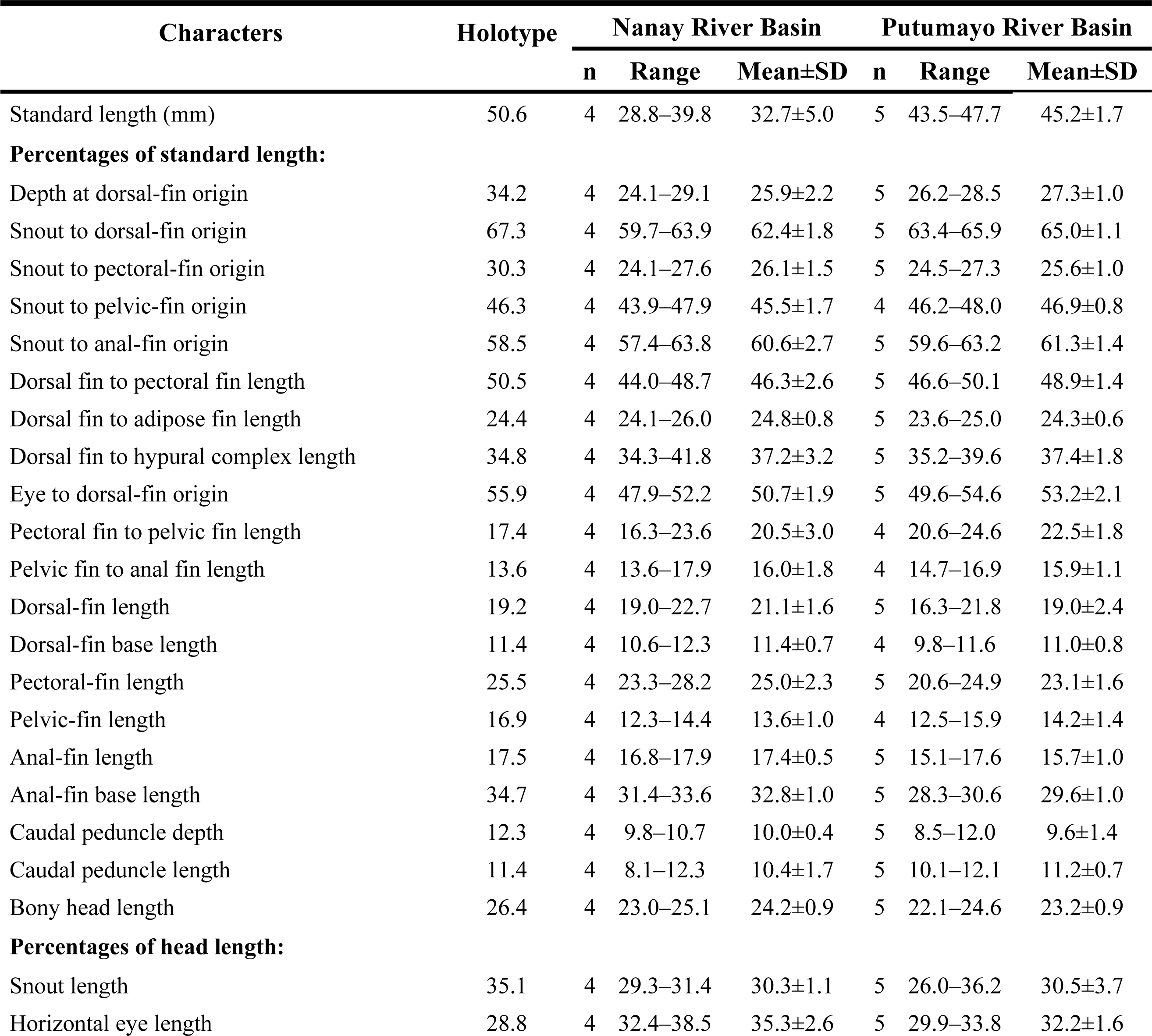
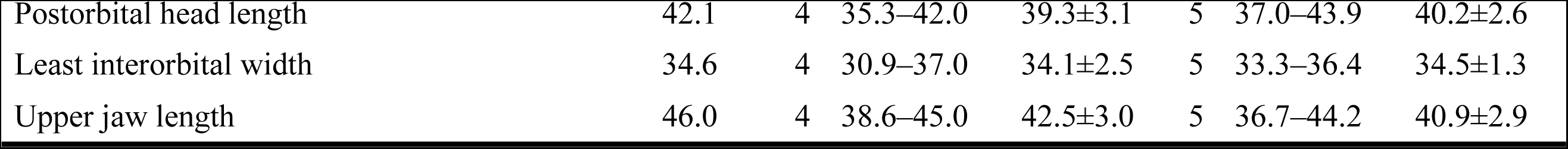
Morphometric data of *Chrysobrycon mojicai* individuals in the Nanay and Putumayo River basins, with information morphometric of holotype (IAvH-P 13932).

Additionally, in our results, Table 2 includes morphometric information of the holotype related to head measurements, which were erroneously digitized in Vanegas et al. [15]: Table 1.

*Chrysobrycon mojicai* is diagnosed by the presence of 10–17 maxillary teeth (12–18 in studied specimens), which are mostly laterally curved distally, dark pigmentation confined to the ventral region between the pelvic and anal fins in adult males (observed in a studied specimens from the Putumayo basin), and 20–27 dentary teeth (15–20 in studied specimens) (Fig 2). The studied specimens were identified as *Chrysobrycon* by having upper mouth, dorsal-fin origin located posterior to anal-fin origin, and frontals extensively contacting along the midline, among other characteristics not associated with the sexual dimorphism. They are recognized as *C. mojicai* by having the distally curved maxillary teeth and similar body shape measurements. We observed that the studied specimens from the Nanay and Putumayo River basins have a fewer number of dentary teeth in comparison with the known range for the species (15–20 *vs.* 20–27).

**Fig 2.**
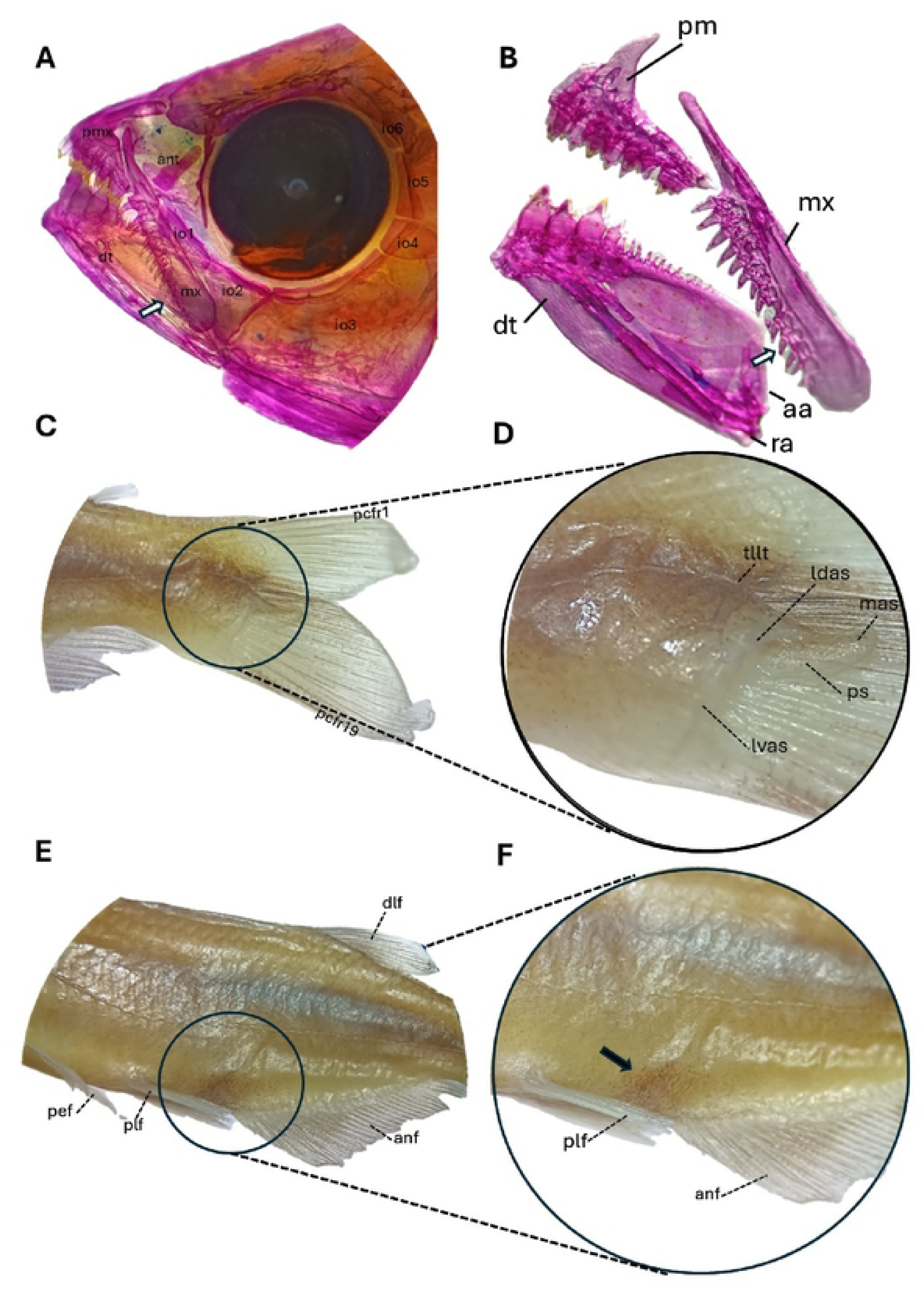
Jaw and dentition, caudal fin, and major sexually dimorphic characteristics of *Chysobrycon mojicai.* (A) Detail of lateral right-side view of the maxilla, premaxilla and dentary in an alcohol-preserved specimen, female, CIIAP 2626, 32.78 mm, from Nanay River basin, (white arrow): curved maxillary teeth. (B) Jaw and dentition of *C. mojicai*, CIIAP 2626, 32.78 mm SL, from Nanay River. (C) Caudal fin of *C. mojicai*, male, CIIAP 2315, 45.89 mm SL, from Putumayo River basin, (D) Details of the hypertrophied caudal-fin squamation on the lower caudal-fin lobe, male, CIIAP 2315, 47.67 mm SL, from Putumayo River basin. (E) Left-side view of body, male, CIIAP 2315, 47.67 mm SL, from Putumayo River basin. (F) Left-side view of distinctive pigmentation (black arrow): around urogenital in adult males, CIIAP 2315, 47.67 mm SL, from Putumayo River basin. Pmx, premaxilla; mx, maxilla; dt, dentary; io, infraorbital; aa, anguloaerticular; ra, retro-articular; pcfr, principal caudal-fin ray; pet, pectoral fin; plf, pelvic fin; and, anal fin; dlf, dorsal fin; lvas, lateroventral accessory scale; ldas, laterodorsal accessory scale; mas, medial accessory scale; ps, pouche scale; pcr,; tllt, terminal lateral-tube.

The discovery of *C. mojicai* from Peruvian waters adds new locality records and helps to elucidate the distribution of this Stevardinii fish in the Amazon basin.

Previously, records of this species were limited to southeastern Colombia [15], which may be partly an artifact of inadequate sampling and taxonomic expertise across the Peruvian region. Notably, our new record represents the first validated occurrence of *C. mojicai* in Peruvian waters (Fig 3).

**Fig 3.**
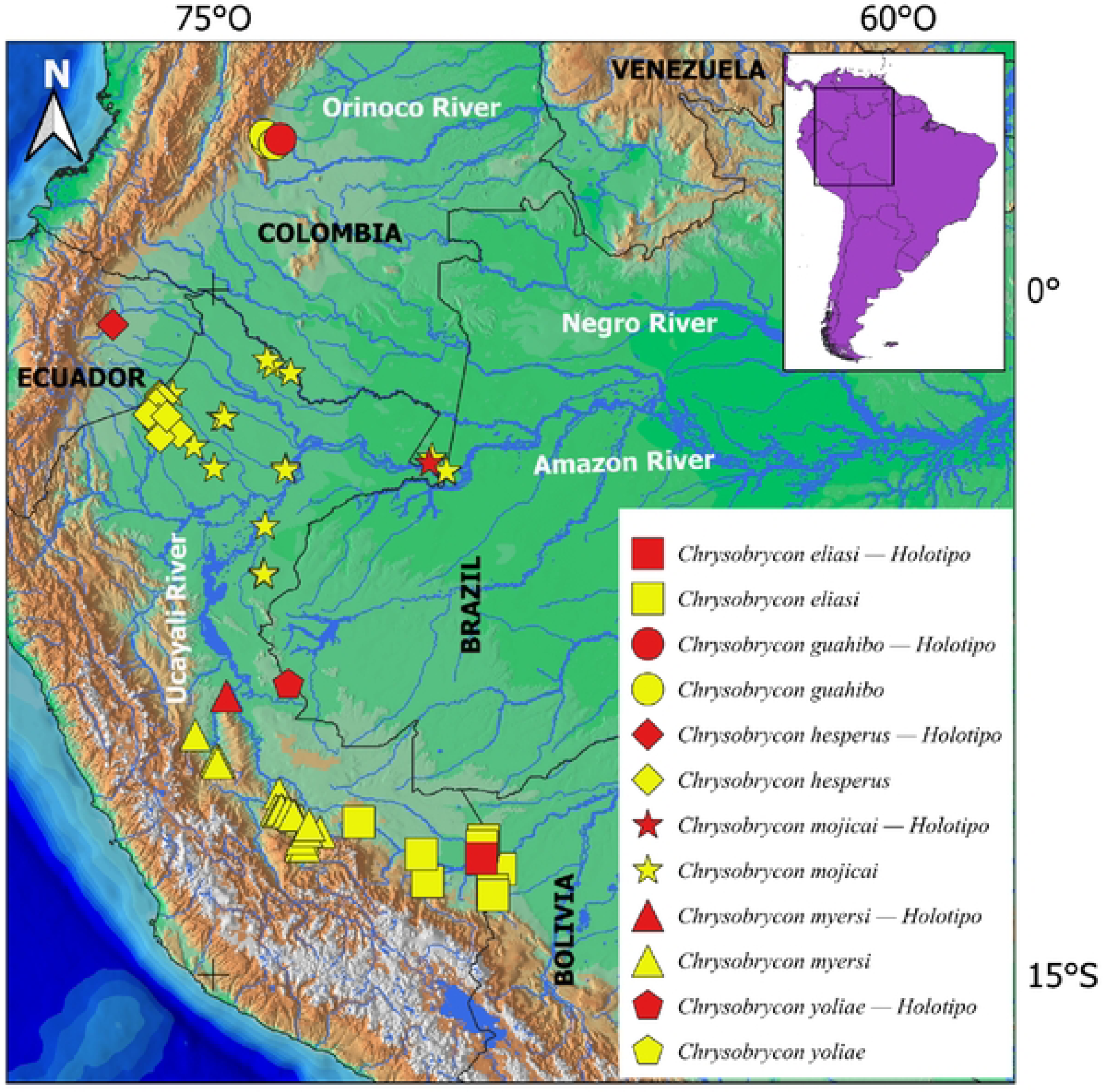
Geographic distribution of *Chrysobrycon* along the mains basin of Peru. Each symbol may represent more than one specimen. Red symbol represents holotype of each species.

### Morphometric analysis

The size-corrected PCA explained 63.9 % of the total variance along the first four components (Table 3), which were identified as the most biologically informative following the scree plot method. The individuals from the Nanay and Putumayo River basins were not separate from the type specimens along PC1 vs. PC2 and PC2 vs. PC3 plots. Along PC1, the female individuals were more strongly discriminated from the male individuals (Fig 4a). The measurements that most strongly influenced PC1 were the depth at dorsal-fin origin (0.7), anal-fin base length (0.6), snout to anal-fin origin (- 0.6), and distance between pectoral- and pelvic-fin origins (-0.5) (Table 4). Due to this sexual differentiation, the specimens from the Nanay and Putumayo basins were more closely grouped with the females paratypes than with the males (Fig 4b).

**Fig 4.**
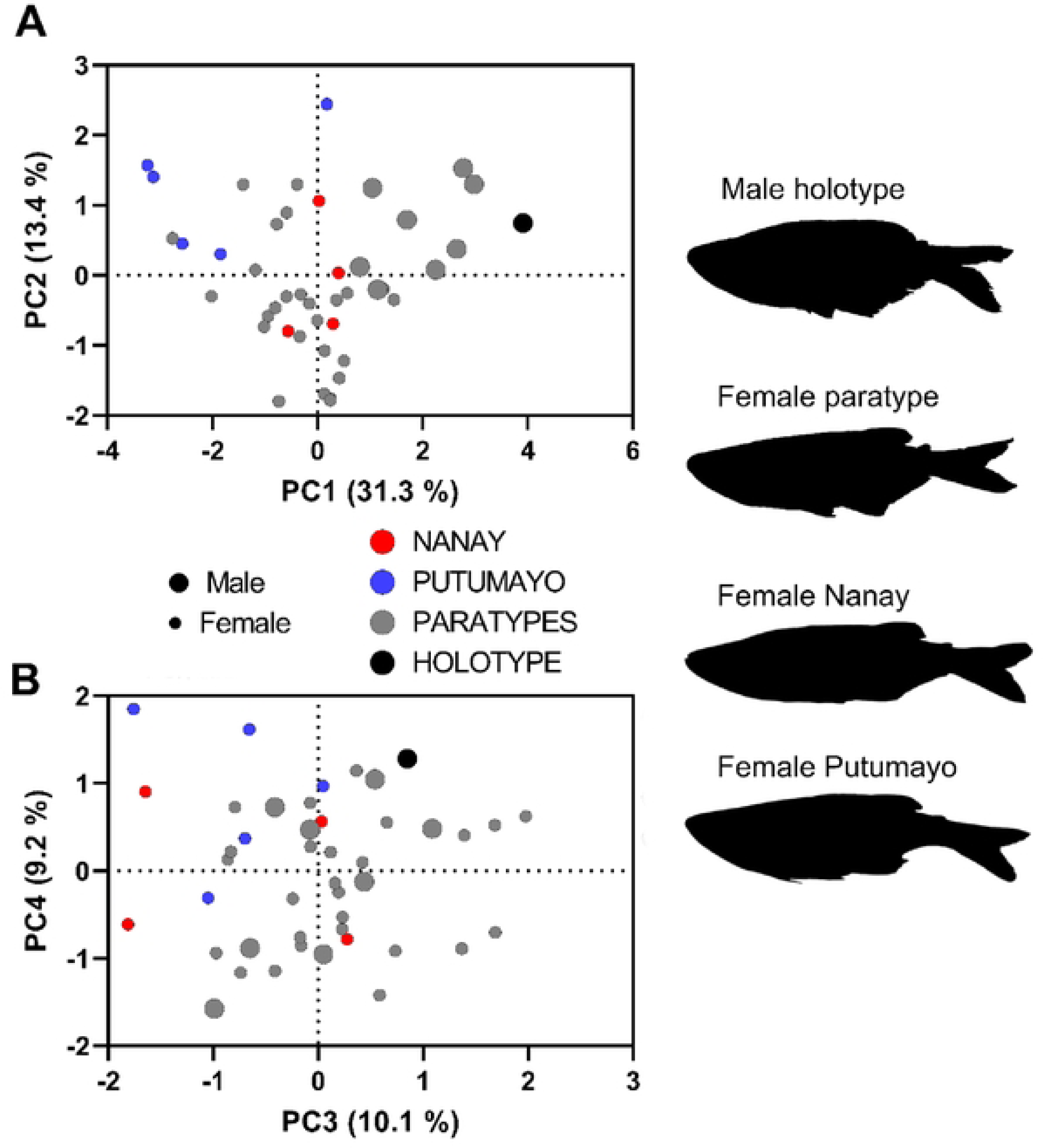
Size-corrected PCA analysis comparing the analyzed specimens of *Chrysobrycon mojicai.* (A). PC1 *vs*. PC2 plot. (B). PC3 *vs.* PC4 plot. Fish silhouette of male holotype, female paratype, and new records here studied are depicted.

**Table 3.** Total variance explained from the size-corrected PCA of the studied specimens of *Chrysobrycon mojicai*.

| PC summary | Eigenvalue | Proportion of variance (%) | Cumulative proportion of variance (%) |
| --- | --- | --- | --- |
| PC1 | 2.5 | 31.3 | 31.3 |
| PC2 | 1.0 | 13.4 | 44.6 |
| PC3 | 0.8 | 10.1 | 54.8 |
| PC4 | 0.7 | 9.2 | 63.9 |
| PC5 | 0.5 | 6.3 | 70.2 |
| PC6 | 0.4 | 4.9 | 75.1 |
| PC7 | 0.3 | 4.5 | 79.6 |
| PC8 | 0.3 | 3.6 | 83.2 |
| PC9 | 0.2 | 2.7 | 85.9 |
| PC10 | 0.2 | 2.5 | 88.3 |
| PC11 | 0.2 | 2.2 | 90.5 |
| PC12 | 0.2 | 2.0 | 92.5 |
| PC13 | 0.1 | 1.6 | 94.1 |
| PC14 | 0.1 | 1.3 | 95.3 |
| PC15 | 0.1 | 1.0 | 96.3 |
| PC16 | 0.1 | 0.9 | 97.1 |
| PC17 | 0.1 | 0.7 | 97.8 |
| PC18 | 0.0 | 0.6 | 98.4 |
| PC19 | 0.0 | 0.5 | 98.9 |
| PC20 | 0.0 | 0.4 | 99.3 |
| PC21 | 0.0 | 0.2 | 99.5 |
| PC22 | 0.0 | 0.2 | 99.7 |
| PC23 | 0.0 | 0.1 | 99.8 |
| PC24 | 0.0 | 0.1 | 100.0 |
| PC25 | 0.0 | 0.1 | 100.0 |
| PC26 | 0.0 | 0.0 | 100.0 |

**Table 4.**
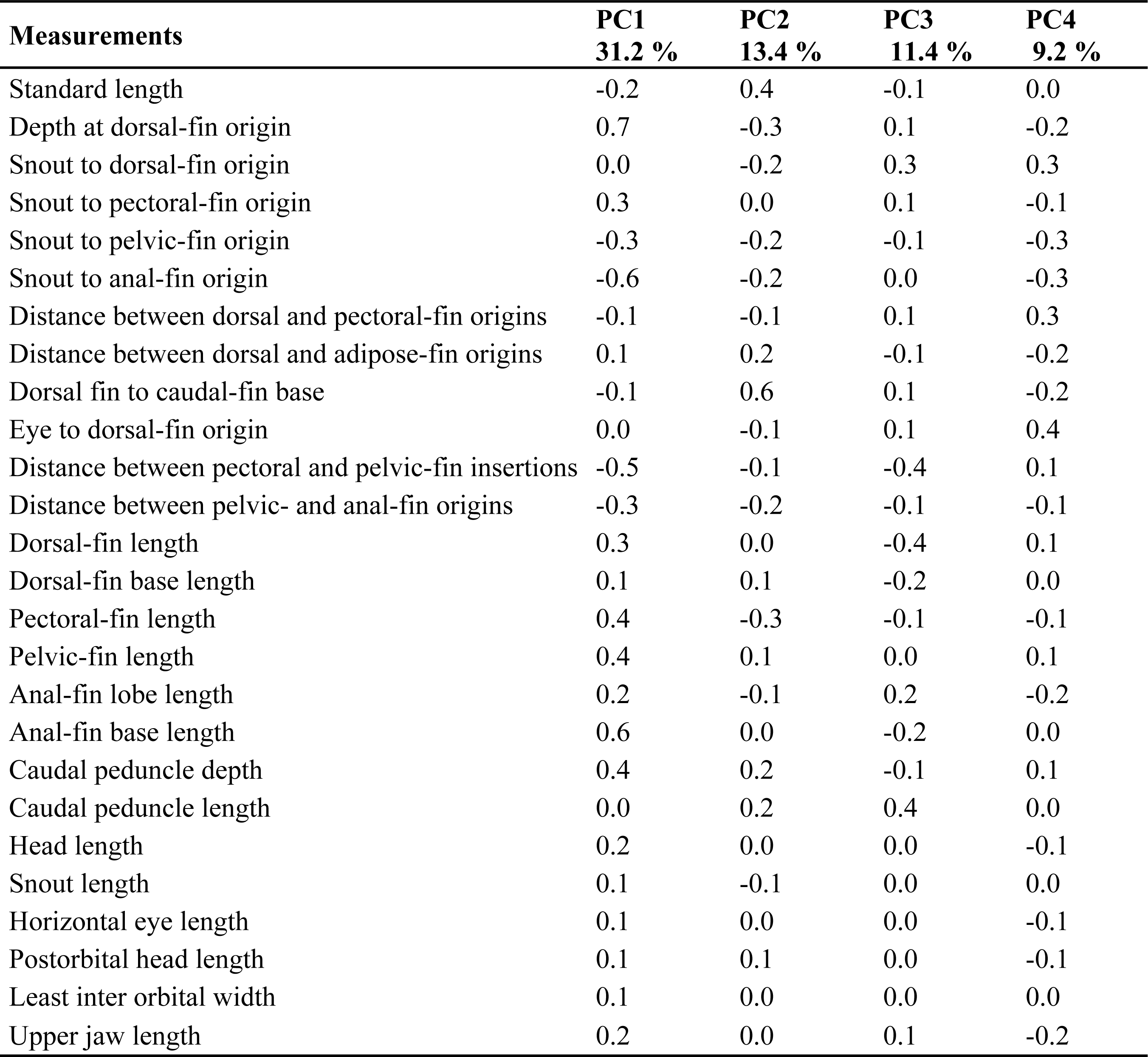
Loadings obtained from the selected components in the size-corrected PCA comparing the studied specimens of *Chrysobrycon mojicai*.

| Measurements | PC1<br>31.2 % | PC2<br>13.4 % | PC3<br>11.4 % | PC4<br>9.2 % |
| --- | --- | --- | --- | --- |
| Standard length | -0.2 | 0.4 | -0.1 | 0.0 |
| Depth at dorsal-fin origin | 0.7 | -0.3 | 0.1 | -0.2 |
| Snout to dorsal-fin origin | 0.0 | -0.2 | 0.3 | 0.3 |
| Snout to pectoral-fin origin | 0.3 | 0.0 | 0.1 | -0.1 |
| Snout to pelvic-fin origin | -0.3 | -0.2 | -0.1 | -0.3 |
| Snout to anal-fin origin | -0.6 | -0.2 | 0.0 | -0.3 |
| Distance between dorsal and pectoral-fin origins | -0.1 | -0.1 | 0.1 | 0.3 |
| Distance between dorsal and adipose-fin origins | 0.1 | 0.2 | -0.1 | -0.2 |
| Dorsal fin to caudal-fin base | -0.1 | 0.6 | 0.1 | -0.2 |
| Eye to dorsal-fin origin | 0.0 | -0.1 | 0.1 | 0.4 |
| Distance between pectoral and pelvic-fin insertions | -0.5 | -0.1 | -0.4 | 0.1 |
| Distance between pelvic- and anal-fin origins | -0.3 | -0.2 | -0.1 | -0.1 |
| Dorsal-fin length | 0.3 | 0.0 | -0.4 | 0.1 |
| Dorsal-fin base length | 0.1 | 0.1 | -0.2 | 0.0 |
| Pectoral-fin length | 0.4 | -0.3 | -0.1 | -0.1 |
| Pelvic-fin length | 0.4 | 0.1 | 0.0 | 0.1 |
| Anal-fin lobe length | 0.2 | -0.1 | 0.2 | -0.2 |
| Anal-fin base length | 0.6 | 0.0 | -0.2 | 0.0 |
| Caudal peduncle depth | 0.4 | 0.2 | -0.1 | 0.1 |
| Caudal peduncle length | 0.0 | 0.2 | 0.4 | 0.0 |
| Head length | 0.2 | 0.0 | 0.0 | -0.1 |
| Snout length | 0.1 | -0.1 | 0.0 | 0.0 |
| Horizontal eye length | 0.1 | 0.0 | 0.0 | -0.1 |
| Postorbital head length | 0.1 | 0.1 | 0.0 | -0.1 |
| Least inter orbital width | 0.1 | 0.0 | 0.0 | 0.0 |
| Upper jaw length | 0.2 | 0.0 | 0.1 | -0.2 |

### Molecular analysis

The sequence alignment of the COI gene yielded 530 base pairs of nucleotides. Based on the percentage pairwise distances obtained, the specimens from the Nanay and Putumayo River basins were separated by 4.4 % (S2 Table 2). The ML tree (Fig 5) did not recover *Chrysobrycon* and *Gephyrocharax* as monophyletic groups. *Chrysobrycon mojicai* was closely related to one specimen of *C. myersi* from the Pachitea River basin (type locality), while other specimens of the latter are associated with specimens identified as *Gephyrocharax* from the Amazon basin, which includes *Gephyrocharax major*. Additionally, a group of *Gephyrocharax* from the trans-Andean region was obtained as monophyletic, but with unclear interspecific relationships. Other genera, such as *Pseudocorynopoma* and *Corynopoma* formed well-defined clades. *Pterobrycon* was recovered as the sister group to *Gephyrocharax* trans-Andean, Sequences from *Hysteronotus* specimens were grouped with Cheirodontinae (S3 Fig 1), demonstrating a misidentification in these sequences. Our results highlight the unresolved taxonomy of the voucher specimens associated with most COI sequences available in GenBank and BOLD of the genera *Chrysobrycon, Gephyrocharax,* and *Hysteronotus*.

**Fig 5.**
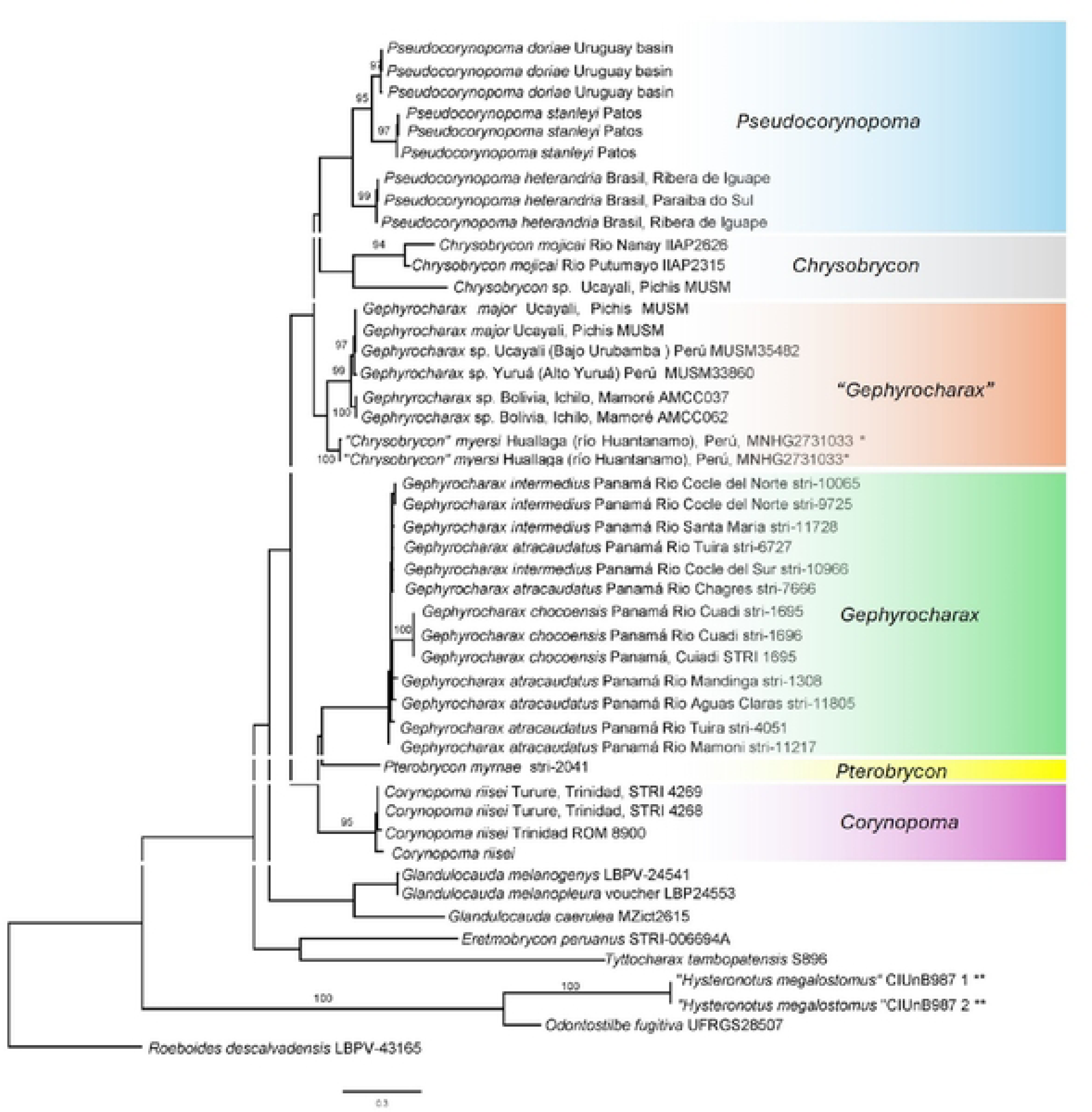
Maximum likelihood topology, using COI data from the Stevardiini tribe. Bootstrap values above 95 % on tree branches.

### Ecological notes

Specimens of *Chrysobrycon mojicai* were collected in blackwater streams of the Nanay River and mixed water streams of the Putumayo River (Fig 6). The habitats where they were captured are highly dependent on precipitation, as the water levels are unstable.

**Fig 6.**
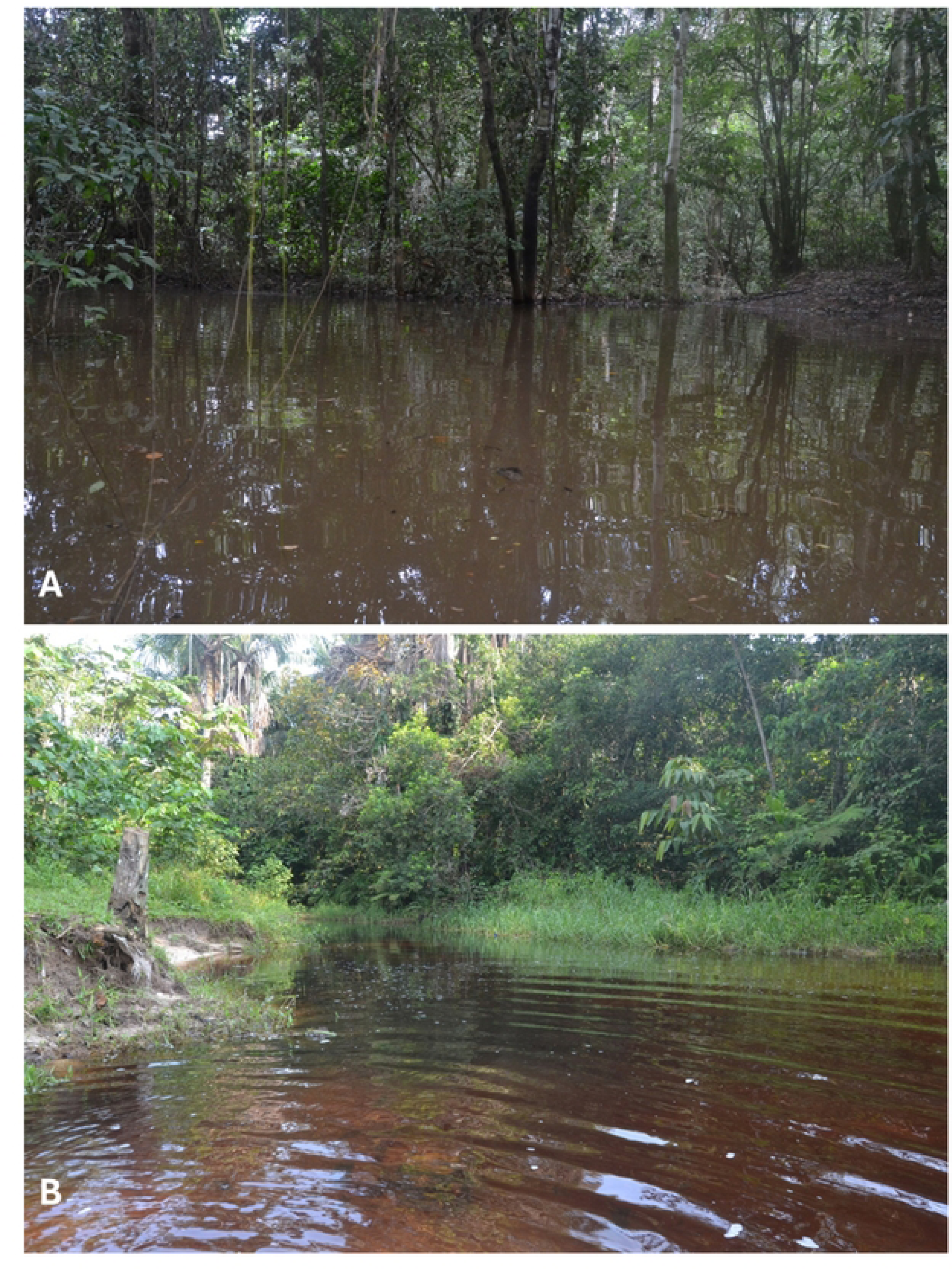
Collection location in the Putumayo and Nanay river basins of *Chrysobrycon mojicai*. (A) Santa Elena creek, Putumayo River basin. (B) San Juan Baustista district, Tanshi creek. Showing the presence of blackwater.

The substrate was sandy, with a slight presence of vegetal material. In the Tanshi stream, the average width was 4.7 meters, and the depth was 0.57 meters. The average physical and chemical parameters of the water were: water temperature 25.41 °C, pH 4.81, dissolved oxygen 1.4 mg/l, electrical conductivity 13 µS/cm, and total suspended solids 7.0 mg/l. For the Santa Elena stream, the average width was 6.1 meters, and the depth was 1.60 meters. The average values of the parameters were: water temperature 25.7°C, pH 7.4, electrical conductivity 100.5 µS/cm, and total suspended solids 68.7 mg/l. *Chrysobrycon mojicai* co-occurs with *Moenkhausia comma* Eigenmann 1908, *M. oligolepis* (Günther 1864), *M. intermedia* Eigenmann 1908, *Bryconops inpai* Knöppel, Junk & Géry 1968, *B. melanurus* (Bloch 1794), *Ctenobrycon hauxwellianus* (Cope 1870), and *Knodus smithi* (Fowler 1913) in tributary streams of the Nanay River, while in the Santa Elena stream, it co-occurs with *Moenkhausia oligolepis*, *Knodus smithi*, and *Astyanax bimaculatus* (Linnaeus 1758).

### Conservation Status

According to the IUCN database, four *Chrysobrycon* species are categorized as follows: *C. hespersus* and *C. myersi* as Least Concern (LC), *C. guahibo* as Vulnerable (VU), and *C. eliasi* as Data Deficient (DD). *C. calamar* has not yet been categorized (NE). The presence of *C. mojicai* in the Peruvian Amazon expands its occurrence to exceed 20,000 km2. Although this species appears to be rare in the Peruvian Amazon, based on its updated distribution range, it is categorized as Least Concern (LC).

## Discussion

The record of *Chrysobrycon mojicai* for the Peruvian Amazon extends its geographic range approximately 400 km northwest to Putumayo River and about 340 km southwest to Nanay River from its type locality. The studied specimens from the Nanay and Putumayo River basins were identified as *C. mojicai* based on morphological, morphometric, and meristic evidence observed in adult females from Nanay River and both males and female from the Putumayo River. Following these analyses, all specimens identified as *Chrysobrycon* sp. in the MUSM collection were reviewed, which allowed us to further extend the distribution of this species to the Tigre and Tapiche River basins. We could not observe some sexual characteristics of the specimens collected from the Nanay River basin because adult males were not collected. However, these characteristics were observed in the individuals from Putumayo River basin (Fig 2). The specimens collected in the Nanay River basin were identified as *C. mojicai* mainly due to the presence of laterally curved maxillary teeth, an exclusive characteristic defining the species. The specimens collected in the Putumayo River basin, which included mature males, presented additional characteristics such as modified caudal fin scales and dark pigmentation located between the pelvic- and anal-fin origins, which is more intense ventrally around the urogenital pore. We obtained an unusual count of dentary teeth in the studied specimens (15–20), which is lower than the known range for the species (20–27) [15]. However, we hope to better understand this discrepancy when more males from the populations analyzed here become available. Based on the maximum likelihood topology using COI, the assignment of the studied specimens is partially attributable to *Chrysobrycon*, given that there is no COI data available for the type specimens of *C. mojicai* and the genus was not recovered as monophyletic.

Meza-Vargas et al. [18] reported *Chrysobrycon guahibo* in the Loreto region, Peru, based on specimens deposited in the MUSM collection. During this study, these individuals were reviewed through morphological analyses and re-evaluated using the taxonomic key of Vanegas-Ríos et al. [14], leading to a correction in their taxonomic identification. The specimens were identified as an undescribed species, *Chrysobrycon* sp. Based on this review, the distribution of *C. guahibo* has been corrected, removing its record from the Peruvian Amazon and maintaining its known distribution in the Orinoco basin. It is important to note that the geographic distribution of *Chrysobrycon* species is not yet fully understood in Bolivia, Colombia, Peru, and Ecuador, despite ongoing study and review of this group in recent years [12, 15, 44].

The genetic distances obtained for *Chrysobrycon* species revealed that specimens from the Nanay and Putumayo basins of *C. mojicai* were differentiated by 4.4%. In contrast, the genetic distance between these specimens and those of *C. myersi* from the Pachitea River basin (type locality) ranged between 11.2% and 12.8%. This pattern, where populations are morphologically similar but exhibit relatively high genetic distances, has also been observed in other fish species. For instance, *Micromyzon akamai* (Friel & Lundberg, 1996) exhibited a genetic distance of 4.0% between populations from the Napo River basin in Ecuador and the lower Amazon basin in Brazil, which is comparable to the 4.0% intraspecific distance reported for *Ernstichthys megistus* (Orcés V., 1961) [45].

Conversely, some fish species exhibit low genetic distances yet have well-marked morphological differences. For example, *Semaprochilodus insignis* (Jardine, 1841) and *S. kneri* (Pellegrin, 1909) have genetic distances of less than 0.5% [46], and *Prochilodus nigricans* (Spix & Agassiz, 1829) and *P. rubrotaeniatus* (Jardine, 1841) show genetic distances of less than 0.3% [47]. Another example is when populations are morphologically distinct but genetically similar, as observed in the stevardiine *Diapoma pampeana*, which has a genetic distance ranging between 0–0.2% [48]. Although the genetic distance between individuals collected from the Nanay and Putumayo Rivers could be considered relatively high for intraspecific variation, the morphological, morphometric, and meristic analyses support their identification as *C. mojicai*. It is important to note that the genetic boundaries within the Stevardiini are not yet fully understood, representing an intriguing area for future research.

In addition to the sequences here obtained for *C. mojicai*, the sequence of *C. myersi* from the Pachitea basin was correctly identified. This situation underscores the scarcity of molecular data available for the genus and its species, highlighting the necessity for further sequencing efforts, particularly for species with type localities in Perú. The phylogenetic tree based on COI data also revealed several unresolved issues within Stevardiini, such as the non-monophyly of certain genera, which extend beyond the scope of this study.

Recent phylogenetic research combining molecular and morphological data has supported the monophyly of Stevardiini, including *Hysteronotus* [9], aligning with earlier morphological studies [10, 49]. However, it is noteworthy that both Thomaz et al. [8] and Ferreira et al. [9] did not include molecular data for *Hysteronotus*. In this study, we incorporated available *Hysteronotus* sequences from GenBank. Our analysis places these *H. megalostomus* samples within the Cheirodontinae clade (Fig. 5 and S3 Fig. 1), suggesting they may belong to a species of *Macropsobrycon*. Morphologically, *H. megalostomus* differs significantly from Cheirodontinae. For example, *H. megalostomus*, recently redescribed by Menezes et al. [50], is characterized by two rows of teeth, the absence of a pseudotympanum, the lack of pedunculated teeth, and the presence of a humeral spot, all of which differentiate it from Cheirodontinae

The non-monophyly of *Chrysobrycon* and *Gephyrocharax* here obtained disagrees with previous phylogenetic analyses [10, 49], but again elucidate the initial findings presented by Thomaz et al. [8] in this regard. The monophyly of *Gephyrocharax* presented by Vanegas-Ríos et al. [10], despite the large morphological dataset, could not strongly supported by high values of node, and as consequence, it is expected that the genus interrelationships need be further studied. Therefore, we hope that a more stable and understandable point of the phylogeny of these genera can be reached combining morphological and molecular evidence, as is being done in ongoing research by one of the co-authors (JAVR). It is important to note that our results are based on a single molecular marker, which may partially explain the discrepancies with previous phylogenetic studies (e.g., [9]). Additionally, a thorough review of the voucher specimens used for COI sequencing within Stevardiini is necessary, as our findings illustrate potential issues with morphological identification (e.g., *Chrysobrycon myersi* from the Huallaga basin and *Hysteronotus megalostomus* from San Francisco River basin).

In recent years, environmental DNA (eDNA) studies have gained traction as powerful tools for biodiversity monitoring and species identification. However, many of these studies face limitations due to the lack of a comprehensive molecular reference database, which often results in sequence identifications being resolved only to the genus level. This can lead to incomplete or ambiguous species identifications, which hampers our understanding of biodiversity and ecosystem health. The molecular data presented in this study significantly enrich the DNA barcoding database for Peru, providing crucial reference sequences that enhance species identification accuracy. By improving the molecular reference base, this study addresses the gaps in our current knowledge and supports more precise and reliable taxonomic assignments. This contribution is particularly valuable for the Peruvian Amazon, a region known for its high biodiversity and ecological complexity.

### Comparative examined material

*Chrysobrycon eliasi*. **all from Peru**: Madre de Dios department, Tambopata: MUSM 39970, holotype, 34.3 mm SL, Madre de Dios basin, Loboyoc creek, 12°27′07S 69°7′43’’W, 210 m asl. MLP 10831, three paratypes, 33.0–43.5 mm SL (two c&s specimens 33.0–39.9 mm SL), Manuripe River basin, creek at km 50, 12°11′21’’S 69°6′57’’W, 248 m asl.

*Chrysobrycon hesperus*. **all from Ecuador:** USNM 164056, holotype of *Hysteronotus hesperus* Böhlke, 1958, 72.3 mm SL (radiograph), Napo-Pastaza, Río Pucuno, tributary of Río Suno, Pucuno, enters of Suno, 0°46’S 77°12’W, approximately 300-350 m asl ((above sea level). USNM 175124, 1 paratype, 59.1 mm SL (radiograph), Napo- Pastaza, Río Pucuno, tributary of Río Suno, Pucuno, enters of Suno, 3°46’S 17°12’W. **All from Peru: Loreto Department: Tapiche River:** MUSM 24543, 7, 25.44-30.26 mm SL, Requena Province, Requena district, Paranayari creek. **Tigre River:** MUSM 26607, 4, 41.66-69.87 mm SL, Loreto Province, Trompeteros district, Caballo creek. MUSM 26617, 3, 29.11-34.26 mm SL, Loreto Province, Trompeteros district, unnamed creek tributary to Huayuri creek. MUSM 28682, MUSM 28682, 16, 21.91-49.42 mm SL. Loreto Province, Tigre district, San Carlos creek. MUSM 28709, 2, 25.01-25.55 mm SL. Loreto Province, Tigre District, Manchari lagoon tributary to Corrientes River. MUSM 28672, 8, 32.69-36.93 mm SL, Loreto Province, Tigre district, Forestal creek. MUSM 30582, 1, 31.63mm SL, Loreto Province, Trompeteros district, Corrientes River. MUSM 30627, 1, 33.24 mm SL, Loreto Province, Trompeteros district, Macusari River. MUSM 32124, 1, 28.02 mm SL, Loreto Province, Trompeteros district, Palatanoyacu River. MUSM 33159, 41, 20.18-45.55 mm SL, Loreto Province, Trompeteros district, Carmen creek. MUSM 33194, 4, 23.95-26.14 mm SL, Loreto Province, Trompeteros district, Paniyacu creek. **Pastaza River:** MUSM 32884, 1, 17.02 mm SL, Datem del Marañon Province, Andoas district, Capahuari creek. **Itaya River:** MUSM 30439, 1, 23.97 mm SL, Maynas Province, Iquitos, San Lucas creek.

*Chrysobrycon guahibo*. **all from Colombia:** Meta department, Orinoco River basin, Guaviare River basin, Ariari River basin: MPUJ 7160, holotype, 31.9 mm SL, Colombia, Fuente de Oro municipality, Caño Abrote, 3°17′39′′N 73°32′02′′W, 258 m asl. MLP 10829, two c&s specimens, 30.4–31.3 mm SL.

*Chrysobrycon myersi*. **all from Peru: Pachite River: Huanuco Department:** ANSP 112325, two paratypes, 30.1–46.1 mm SL, small creek at northeastern outskirts of Tournavista. ANSP 112326, three paratypes, 28.3–32.0 mm SL, small creek at northeastern outskirts of Tournavista. **Pasco Department:** MUSM 67623, 1, 48.57 mm SL, Oxapampa Province, Puerto Bermudez district, Pichis River. **Urubamba River: Cusco Department:** MUSM 59970, 3, 30.97-48.18 mm SL, La Convencion Province, Megantoni district, Urubamba River. MUSM 68082, 3, 48.46-52.65 mm SL, La Convencion Province, Megantoni district, Camisea River. MUSM 70278, 2, 40.42-45.28 mm SL, La Convencion Province, Megantoni district, unnamed creek tributary to Urubamba River. MUSM 70283, 1, 56.63 mm SL, La Convencion Province, Megantoni district, unnamed creek tributary to Urubamba River. MUSM 70289, 1, 51.44 mm SL, La Convencion Province, Megantoni district, unnamed creek tributary to Urubamba River. MUSM 70292, 2, 45.28-46.07mm SL, La Convencion Province, Megantoni district, unnamed creek tributary to Urubamba River. MUSM 70295, 1, 41.57 mm SL, La Convencion Province, Megantoni district, unnamed creek tributary to Urubamba River. MUSM 70332, 2, 52.61-62.98 mm SL, La Convencion Province, Megantoni district, Huitricaya River. MUSM 71820, 3, 38.52-49.45 mm SL, La Convencion Province, Megantoni district, Huitricaya River. MUSM 71903, 4, 44.01-63.94 mm SL, La Convencion Province, Megantoni district, unnamed creek tributary to Huitricaya River. MUSM 71945, 3, 45.15-46.21 mm SL, La Convencion Province, Megantoni district, Choro creek tributary to Camisea River. **Junin Department:** MUSM 70298, 91, 26.32-56.91 mm SL, Satipo Province, Rio Tambo district, Shotariari creek. MUSM 70301, 54, 28.11-59.89 mm SL, Satipo Province, Rio Tambo district, Shotariari creek. MUSM 70315, 19, 39.48-66.31 mm SL, Satipo Province, Rio Tambo district, Santonaro creek.

*Chrysobrycon mojicai*. **all from Colombia:** Amazonas department, IAvH-P 13932, holotype male, 50.6 mm SL, Leticia, Amacayacu National Natural Park, Amazon River basin, unnamed forest stream tributary of Mata-matá River, 3°48′23′′S 70°15′58′′W, 86 m asl, 25 March 2006, F. Arbeláez & unknown collectors. IAvH-P 8291, five specimens, 25.0–50.4 mm SL (one c&s specimen, 50.4 mm SL), collected with holotype. IAvH-P 8295, nine specimens, 29.0–47.7 mm SL, Leticia, Amacayacu National Natural Park, Amazon River basin, unnamed forest stream tributary of Pureté River headwaters, 3°41′54′′S 70°12’24′′W 127 m asl, 25 March 2006, F. Arbeláez & unknown collectors. IAvH-P 8300, two specimens, 33.5–40.8 mm SL, Leticia, Amacayacu National Natural Park, Amazon River basin, unnamed forest stream tributary of Pureté River headwaters, 3°41′37.5′′S 70°12’26.5′′W, F. Arbeláez & unknown collectors, 26 March 2006. IAvH-P 8917, 14 specimens, 17.1–47.5 mm SL, Leticia, Amazon River basin, Amazon River basin, Sufragio stream in front of Zafire Biological Station, 4°0′19.2′′S 69° 53′55.5′′W 118 m asl, 8 December 2005, F. Arbeláez. IAvH-P 8951, nine specimens, 17.9–50.5 mm SL, Sufragio stream in front of Zafire Biological Station, 4°0′19.2′′S 69° 53′55.5′′W 118 m asl, 9 December 2005, F. Arbeláez. IAvH-P 9022, six specimens, 43.8–50.8 mm SL, Leticia, Amazon River basin, Sufragio stream in front of Zafire Biological Station, 4°0′19.2′′ S 69°53′55.5′′ W 118 m asl, 14 December 2005, F. Arbeláez. IAvH-P 9070, four specimens, 48.6–55.0 mm SL, Leticia, Amazon River basin, unnamed forest stream tributary of Calderon River, 45 min. NE of Zafire Biological Station, 3°58′40.1′′S 69°53′31.8′′W 130 m asl, 12 Dec 2005, F. Arbeláez. IAvH-P 9093, four specimens, 23.7–47.8 mm SL, Leticia, Amazon River basin, unnamed stream, tributary of Calderon River, 45 min. NE of Zafire Biological Station, 3°58′40.14′′S 69°53′31.8′′W 130 m asl, 13 December 2005, F. Arbeláez. MPUJ 8058, one specimen, 49.5 mm SL, collected with holotype. MPUJ 8059, one c&s specimen, 50.3 mm SL, Leticia, Amazon River basin, unnamed forest stream tributary of Calderon River, 45 min. NE of Zafire Biological Station, 3°58′40.1′′S 69°53′31.8′′W 130 m asl, 12 Dec 2005, F. Arbeláez.

*Chrysobrycon yoliae*. **all from Peru:** Ucayali department, Coronel Portillo province, Abujao, Yucamia River subsystem, unnamed creek, 8°39′14′′S 73°21′17′′W, approximately 273 m asl: MUSM 46140, holotype, 51.6 mm SL. MLP 10517, one paratype, 48.4 mm SL. MUSM 46141, eight paratypes, 38.2–51.5 mm SL.

## Acknowledgements

The study was carried out with the support of the Institute of Research of the Peruvian Amazon (IIAP), as part of the project named "Binational Expedition Peru-Colombia: Biological Diversity Inventories in the Great Putumayo." Additionally, we express our warm gratitude for the local support from the native community of Bobona, in the province of Putumayo, Loreto region. To Sindy Jean Luca, for her assistance in field sampling. Part of the collection corresponds to the project assessing fish diversity in streams along the Iquitos-Nauta road axis, Loreto, Peru, with field support from Edwin Agurto. We thank the CONCYTEC through the PROCIENCIA program within the framework of the contest "Projects for the incorporation of postdoctoral researchers in Peruvian institutions” to support JC [PE501080808-2022], VMV [074-2021; PE501084299-2023-PROCIENCIA-BM] and DF [PE501085130-2023]. JARV is grateful for the financial support from CONICET and FONCyT (PIBAA 0654CO and BID-PICT 2019–02419).

## Author contributions

## Supporting information

**S1 Table 1.** Sequences of species generated in this study (in bold) and those download from GenBank and BOLD used in the analyses

**S2 Table 2.** Estimates of Evolutionary Divergence over Sequence Pairs between groups of genera. Values refer to the genetic distance based on COI sequences

**S3 Figure 1.** Stevardinii Neighbor joining (NJ) phylogeny tree based on cytochrome c oxidase I (COI) gene sequences generated in MEGA 11

